# User-friendly exploration of epigenomic data in single cells using sincei

**DOI:** 10.1101/2024.07.27.605424

**Authors:** Vivek Bhardwaj, Soufiane Mourragui

**Affiliations:** Institute of Biodynamics and Biocomplexity, Department of Biology, Utrecht University, The Netherlands; Hubrecht Institute-KNAW (Royal Netherlands Academy of Arts and Sciences), Utrecht, The Netherlands; University Medical Center Utrecht, Utrecht, The Netherlands

## Abstract

Emerging single-cell sequencing protocols allow researchers to study multiple layers of epigenetic regulation while resolving tissue heterogeneity. However, despite the rising popularity of such single-cell epigenomics assays, the lack of easy-to-use computational tools that allow flexible quality control and data exploration hinders their broad adoption. We introduce the Single-Cell Informatics (**sincei**) toolkit. sincei provides an easy-to-use, command-line interface for the exploration of data from a wide range of single-cell (epi)genomics protocols directly from aligned reads stored in .bam files. sincei can be installed via bioconda and the documentation is available at https://sincei.readthedocs.io.

## Main

Understanding the control of gene expression requires methods that quantify different layers of epigenetic regulation, such as histone modifications, transcription factor binding sites, chromatin conformation, as well as single nucleotide level changes such as DNA methylation. Over the years, several sequencing methods have been developed to study these events genome-wide from a pool of cells, and challenges have been addressed to process and analyze these data. More recently, these methods have been adapted to reach single-cell resolution, allowing researchers to gain a quantitative understanding of cell-to-cell variability in gene regulation^1^. Although single-cell epigenomic technologies are getting increasingly popular, certain issues related to data analysis severely limit their broad adaptability among researchers. First, almost all the existing tools currently follow the standards of single-cell RNA-seq analysis, despite evidence showing that data from epigenomics protocols require a different preprocessing, quality control (QC), and downstream analysis^2^. Second, most popular existing tools and workflows are focused on droplet-based scRNA and ATAC-seq protocols, even though many existing protocols have already been adapted to study epigenetic processes, such as histone modifications, DNA methylation, and 3D genome organization^3^. Finally, the use of current tools requires significant scripting and bioinformatics experience, creating a barrier to entry for biologists. To address this issue, we have developed the **Sin**gle-**Ce**ll **I**nformatics toolkit (**sincei**). This easy-to-use command-line software extends beyond scRNA-seq or scATAC-seq analysis and enables flexible exploration of chromatin, transcriptome, and methylome data. Sincei adapts bulk epigenomics analysis standards into single-cell workflows and simplifies data integration.

The sincei workflow begins directly from BAM formatted files, output from most genome alignment tools, allowing users to aggregate single-cell signals at different genomic resolutions, or employing user-defined features. Starting from BAM files also allows users to perform extensive and flexible read-level QC, such as asking how many reads in a cell do not meet the desired alignment criteria, map to undesired genomic locations, or have undesired GC content. These choices are instrumental in filtering bad-quality cells and regions while retaining useful signals. Similar to other popular tools^4,5^, sincei allows QC and filtering of cells after signal aggregation into the desired bins/cells, all directly from the command line. To facilitate easy exploration of QC metrics, the output of sincei QC steps can be visualized with the MultiQC tool^6^. Provided pre-defined groups (such as cell clusters, batches, or timepoints), sincei also allows users to aggregate signal as pseudo-bulks, creating coverage files (bigwigs) for visualization in IGV^7^, or plotting via deepTools^8^.

Dimensionality reduction and clustering of cells based on genomic signals is another critical step in single-cell data exploration. Both scRNA-seq and scATAC-seq data are count-based. However, the acosh transformation, followed by principal component analysis (PCA) is the most popular workflow of choice for scRNA-seq^9^, while topic models such as Latent Semantic Analysis (LSA) and Latent Dirichlet Allocation (LDA) are more popular for scATAC-seq^10^. On the other hand, DNA-methylation assays produce ratios of signal and require a different modeling approach^11^. sincei accommodates these popular choices through the log1p-PCA, LSA, and LDA methods. Furthermore, we also offer a scalable implementation of GLM-PCA^12,13^, a flexible dimensionality reduction method that can accommodate a broad range of signal types, such as counts (Poisson), binary (Bernoulli), ratio (Beta). Developers can easily extend this list, providing users with the flexibility needed to explore diverse statistical modeling options, particularly instrumental in developing novel assays. Finally, to complete the exploration workflow, sincei also provides a wrapper for graph-based clustering of the resulting signals and their visualization onto 2D UMAPs^14^.

**Fig 1.**
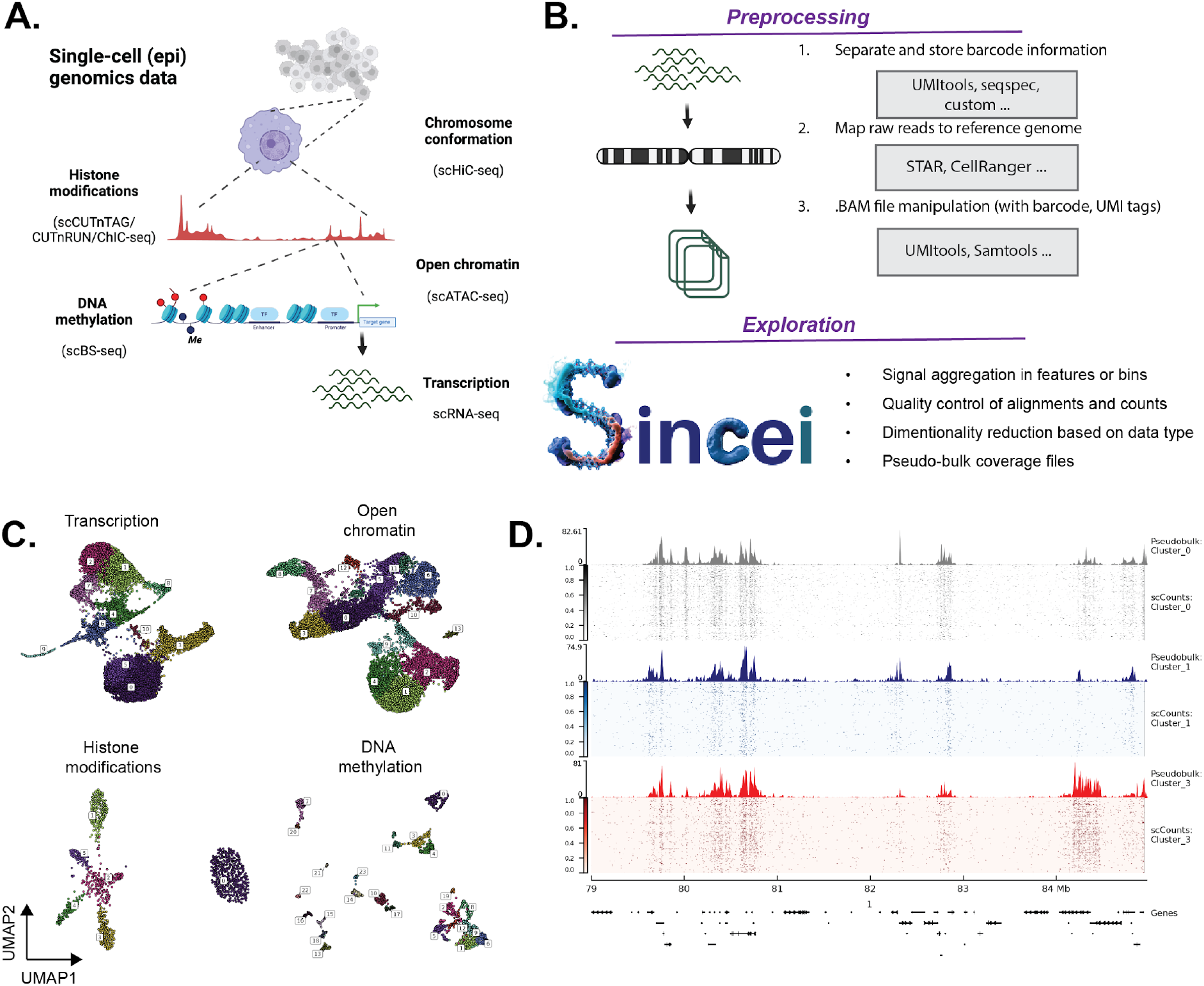
sincei supports exploration workflows for a wide range of single-cell (epi)genomics data. **A**. The range of (epi)genomic protocols that map different levels of gene regulation in single cells (Created with BioRender). **B**. Common steps in the pre-processing workflow for such data can be accomplished using existing tools, while sincei can be used for downstream exploration once the mapped “.bam” formatted files are obtained. **C**. UMAP plots showing 2D projection and clustering results of sincei for 4 different protocols: (top) scRNA-seq and scATAC-seq data from 10x genomics multiome assay of human CD4+ blood cells^15^, (bottom) scSortChIC data mapping H3K4me1 in cells from mouse bone marrow^16^ and snmCAT-seq data mapping CpG methylation in human brain^17^. **D**. pyGenomTracks visualization^18^ of binned counts in single cells, together with the corresponding CPM-normalized pseudo-bulk bigwigs of 3 clusters obtained using sincei on scSortChIC data.

Overall, sincei is a powerful toolkit for “data exploration”. We aim to provide users with tools to efficiently assess the complexity and variance in single-cell signal in their assay. More sophisticated tasks, such as batch effect aware data integration, or gene regulatory network inference would require a deeper dive into the data and are supported by a range of existing toolkits in R or Python^2,19,20^. The command-line tools in sincei ensure easy integration into scalable workflows and make it easier to be used for teaching and training, facilitating the introduction of single-cell genomics data analysis for novice users. For bioinformaticians, our Python API allows a more in-depth exploration of their data. The output files produced by sincei are fully compatible with scverse and bioconductor tools, ensuring interoperability^21,22^. With sincei, we hope to simplify the extraction of biological insights from different genomic layers and facilitate the integration of single-cell and bulk epigenomic assays.

## Code availability

sincei is available open-source at https://github.com/bhardwaj-lab/sincei and can be installed via bioconda (*conda install -c bioconda sincei*). Documentation is available at https://sincei.readthedocs.io.

## Data availability

Data used for sincei documentation and in this manuscript are available from the published studies^15–17^.

## Acknowledgements

This work was supported, in part by the EMBO Long-term Fellowship (ALTF 1197–2019) to V.B. We thank the van Oudenaarden lab at the Hubrecht Institute for providing access to in-house datasets for testing sincei.

## References

1. Vandereyken, K., Sifrim, A., Thienpont, B. & Voet, T. Methods and applications for single-cell and spatial multi-omics. Nat. Rev. Genet. 1–22 (2023).

2. Heumos, L. et al. Best practices for single-cell analysis across modalities. Nat. Rev. Genet. 1–23 (2023).

3. Preissl, S., Gaulton, K. J. & Ren, B. Characterizing cis-regulatory elements using single-cell epigenomics. Nat. Rev. Genet. (2022) doi:10.1038/s41576-022-00509-1.

4. Wolf, F. A., Angerer, P. & Theis, F. J. SCANPY: large-scale single-cell gene expression data analysis. Genome Biol. 19, 15 (2018).

5. Hao, Y. et al. Integrated analysis of multimodal single-cell data. Cell 184, 3573–3587.e29 (2021).

6. Ewels, P., Magnusson, M., Lundin, S. & Käller, M. MultiQC: summarize analysis results for multiple tools and samples in a single report. Bioinformatics 32, 3047–3048 (2016).

7. Robinson, J. T. et al. Integrative genomics viewer. Nat. Biotechnol. 29, 24–26 (2011).

8. Ramírez, F. et al. deepTools2: a next generation web server for deep-sequencing data analysis. Nucleic Acids Res. 44, W160–5 (2016).

9. Ahlmann-Eltze, C. & Huber, W. Comparison of transformations for single-cell RNA-seq data. Nat. Methods 20, 665–672 (2023).

10. Chen, H. et al. Assessment of computational methods for the analysis of single-cell ATAC-seq data. Genome Biol. 20, 241 (2019).

11. Lareau, C., Kangeyan, D. & Aryee, M. J. Preprocessing and Computational Analysis of Single-Cell Epigenomic Datasets. Methods Mol. Biol. 1935, 187–202 (2019).

12. Townes, F. W., Hicks, S. C., Aryee, M. J. & Irizarry, R. A. Feature selection and dimension reduction for single-cell RNA-Seq based on a multinomial model. Genome Biol. 20, 295 (2019).

13. Mourragui, S. et al. Designing DNA-based predictors of drug response using the signal joint with gene expression. (2022).

14. Becht, E. et al. Dimensionality reduction for visualizing single-cell data using UMAP. Nat. Biotechnol. (2018) doi:10.1038/nbt.4314.

15. Persad, S. et al. SEACells infers transcriptional and epigenomic cellular states from single-cell genomics data. Nat. Biotechnol. (2023) doi:10.1038/s41587-023-01716-9.

16. Zeller, P. et al. Single-cell sortChIC identifies hierarchical chromatin dynamics during hematopoiesis. Nat. Genet. (2022) doi:10.1038/s41588-022-01260-3.

17. Luo, C. et al. Single nucleus multi-omics identifies human cortical cell regulatory genome diversity. Cell Genom 2, (2022).

18. Lopez-Delisle, L. et al. pyGenomeTracks: reproducible plots for multivariate genomic datasets. Bioinformatics 37, 422–423 (2021).

19. Kumar, N., Mishra, B., Athar, M. & Mukhtar, S. Inference of Gene Regulatory Network from Single-Cell Transcriptomic Data Using pySCENIC. Methods Mol. Biol. 2328, 171–182 (2021).

20. Kamimoto, K. et al. Dissecting cell identity via network inference and in silico gene perturbation. Nature 614, 742–751 (2023).

21. Virshup, I. et al. The scverse project provides a computational ecosystem for single-cell omics data analysis. Nat. Biotechnol. 41, 604–606 (2023).

22. Amezquita, R. A. et al. Orchestrating single-cell analysis with Bioconductor. Nat. Methods 17, 137–145 (2020).

